# A conserved lysine/arginine-rich motif in potyviral 6K1 protein is key in engaging autophagy-mediated self-degradation for completing pepper veinal mottle virus infection

**DOI:** 10.1101/2024.04.22.590661

**Authors:** Weiyao Hu, Changhui Deng, Li Qin, Peilan Liu, Linxi Wang, Xiaoqin Wang, Wei Shi, Asma Aziz, Fangfang Li, Xiaofei Cheng, Aiming Wang, Zhaoji Dai, Xiaohua Xiang, Hongguang Cui

**Affiliations:** Key Laboratory of Green Prevention and Control of Tropical Plant Diseases and Pests (Ministry of Education) and School of Tropical Agriculture and Forestry, Hainan University, Haikou, China; State Key Laboratory for Biology of Plant Diseases and Insect Pests, Institute of Plant Protection, Chinese Academy of Agricultural Sciences, Beijing, China; College of Plant Protection/Key Laboratory of Germplasm Enhancement, Physiology and Ecology of Food Crops in Cold Region of Chinese Education Ministry, Northeast Agricultural University, Harbin, China; London Research and Development Centre, Agriculture and Agri-Food Canada, London, Ontario, Canada; Haikou Cigar Research Institute, Hainan Provincial Branch of China National Tobacco Corporation, Haikou, China

**Keywords:** *Potyvirus*, autophagy, 6K1, degradation, point mutation, pro-viral role

## Abstract

Potyviruses possess one positive-sense single-stranded RNA genome mainly with polyprotein processing as their gene expression strategy. The resulting polyproteins are proteolytically processed by three virus-encoded proteases into 11 or 12 mature proteins. One of such, 6-kDa peptide 1 (6K1), is an understudied viral factor. Its function in viral infection remains largely mysterious. This study is to reveal part of its roles by using pepper veinal mottle virus (PVMV) as a model virus. Alanine substitution screening analysis revealed that 15 out of 17 conserved residues across potyviral 6K1 sequences are essential for PVMV infection. However, 6K1 protein is less accumulated in virus-infected cells, even though P3-6K1 junction is efficiently processed by NIa-Pro for its release, indicating that 6K1 undergoes a self-degradation event. Mutating the cleavage site to prevent NIa-Pro processing abolishes viral infection, suggesting that the generation of 6K1 along with its degradation might be important for viral multiplication. We corroborated that cellular autophagy is engaged in 6K1’s degradation. Individual engineering of the 15 6K1 variants into PVMV was performed to allow for their expression along with viral infection. Five of such variants, D30A, V32A, K34A, L36A, and L39A, significantly interfere with viral infection. The five residues are enclosed in a conserved lysine/arginine-rich motif; four of them appear to be crucial in engaging autophagy-mediated self-degradation. Based on these data, we envisaged a scenario that potyviral 6K1s interact with an unknown anti-viral component to be co-degraded by autophagy to promote viral infection.

**IMPORTANCE:** *Potyvirus* is the largest genus of plant-infecting RNA viruses, which encompasses socio-economically important virus species, such as *Potato virus Y*, *Plum pox virus*, and *Soybean mosaic virus*. Like all picorna-like viruses, potyviruses express their factors mainly via polyprotein processing. Theoretically, viral factors P3 through CP, including 6K1, should share an equivalent number of molecules. The 6K1 is small in size (∼6 kDa) and conserved across potyviruses, but less accumulated in virus-infected cells. This study demonstrates that cellular autophagy is engaged in the degradation of 6K1 to promote viral infection. In particular, we found a conserved lysine/arginine-rich motif in 6K1s across potyviruses that is engaged in this degradation event. This finding reveals one facet of a small protein that help understand the pro-viral role of cellular autophagy in viral infection.

## INTRODUCTION

Plant viruses are characterized by small genome sizes and compact structures. To overcome their limited coding capacity, viruses have evolved varied gene expression strategies to generate more functional units that are engaged in replication, encapsidation, movement, counter-defense, and transmission. *Potyvirus* is the largest genus of RNA viruses in the plant kingdom, including many well-known viral agents that adversely affect agriculturally and economically important crops, such as potato virus Y (PVY), plum pox virus (PPV), soybean mosaic virus (SMV), and turnip mosaic virus (TuMV) (1–6). All potyviruses possess one single-stranded, positive-sense RNA genome (∼9.7 kb) with a viral protein genome-linked (VPg) covalently linked to its 5′ end and a poly(A) tail at the 3′ terminus, which contains a long, full-genome open reading frame (ORF) and another relatively short ORF (PIPO) embedded in P3-coding region (7, 8). PIPO becomes translational in frame with the coding region of P1 through P3 N-terminus (P3N) from viral genomic subpopulation, which are originated from viral RNA polymerase slippage at a conserved G_1-2_A_6_ motif between *P3N* and PIPO during viral replication (9–11). A similar slippage event occurs in the P1-coding region for sweet potato-infecting potyviruses, giving rise to one more translational ORF (PISPO) in frame with the coding sequence of P1 N-terminus (10, 12, 13). Upon translation, the resulting polyproteins are proteolytically processed by three virus-encoded protease domains (P1, HCPro, and NIa-Pro) into 11 or 12 mature viral units, including two smallest proteins, 6-kDa peptide 1 (6K1) and 6-kDa peptide 2 (6K2) (1, 6). Intriguingly, a recent report showed that potyviral antisense genomes encode small peptides that seem to be essential for viral infectivity (14–16).

The majority of potyviral factors have been substantially studied, and the readers are referred to several recent excellent reviews that summarize their functions in viral infection (3, 4, 6, 17, 18). Potyviral 6K2 is an integral membrane protein and induces endoplasmic reticulum (ER)-derived replication vesicles that move to chloroplast for robust viral replication (19–23). In contrast, the functional roles of the other small peptide, 6K1, are less understood. The 6K1 protein was first defined over 3 decades ago, along with *in vitro* characterization of NIa-Pro cleavage sites at P3-6K1 and 6K1-CI junctions (24, 25). The proteolytic processing at 6K1-CI junction by NIa-Pro is efficient, whereas the cleavage between P3 and 6K1 is slow, when tested in *in vitro* assays or insect cells (25–27). Thus, it was proposed that not only the mature P3 and 6K1, but also the intermediate precursor P3-6K1, are generated from corresponding genomic region (26). As anticipated, both P3 and the precursor P3-6K1 were immuno-detected in tobacco vein mottling virus-infected tobacco leaves and protoplasts by using a polyclonal antibody against P3-6K1 (24). The 6K1 as a mature protein was first detected in PPV-infected *Nicotiana benthamiana* plants via affinity-purified enrichment followed by immuno-detecting using 6K1-specific polyclonal antiserum (28). The 6K1 sequence seems pivotal for viral multiplication. Kekarainen and colleagues adopted transposition-based *in vitro* insertional mutagenesis strategy to generate a genomic 15-bp insertion mutant library based on potato virus A (PVA), and demonstrated that four insertions in the 5’-terminus of *6K1* cistron compromised viral replication (29). In addition, individual deletion of four different motifs in PPV 6K1 abolish viral replication (30). For tobacco vein banding mosaic virus, the mutations introduced into a conserved RSD motif in the middle region of 6K1 inhibited viral replication (31). Importantly, PPV 6K1 is required for viral replication and forms punctate inclusions that target 6K2-induced viral replication complex (VRC) at the early stage of infection (30). In addition, a recent report revealed a counter-defense role of PVY 6K1 via interfering with the interaction of 14-3-3h and TCTP in *N. benthamiana* (32). Nevertheless, a comprehensive investigation on the expression of 6K1 during viral infection and its biological relevance is needed.

Autophagy is an evolutionarily conserved intracellular degradation pathway, by which the damaged or unwanted intracellular components are engulfed by *de novo*-formed double-membrane vesicles (termed autophagosomes) and subsequently delivered to vacuoles for breakdown and turnover in plants (33–35). Numerous recent studies demonstrate that autophagy is engaged in plant defense responses against viruses (including potyviruses), and in turn, viruses evolve strategies to counteract, manipulate or hijack autophagy pathway to promote viral infection (36–38). Potyviral HCPro functions as viral suppressor of RNA silencing (VSR), mainly via directly interacting with, and kidnapping, virus-derived small interfering RNAs (vsiRNAs) (39). Tobacco calmodulin-like protein (rgs-CaM) targets dsRNA-binding domain in HCPro and cooperates with autophagy pathway to degrade HCPro (40). NBR1, a canonical cargo receptor in selective autophagy, targets TuMV HCPro-induced RNA granules for autophagic degradation to suppress viral accumulation (41). Beclin1/ATG6, a core component of phosphoinositide-3-kinase (PI3K) complex, interacts with TuMV NIb and mediates its autophagic degradation likely through an adaptor ATG8a (42).

However, several studies revealed the pro-viral roles of autophagy in potyviral infection. VPg is another potyvirus-encoded VSR, which functions through interacting with suppressor of gene silencing 3 (SGS3, a core component in dsRNA synthesis) to mediate the degradation of SGS3 and RNA-dependent RNA polymerase 6 (RDR6) via both the ubiquitin-proteasome and autophagy pathways (43). Group 1 Remorins (REMs) negatively regulate the cell-to-cell movement of TuMV. To survive, virus-encoded VPg interacts with REM1.2 to degrade it via both 26S ubiquitin–proteasome and autophagy pathways (44). TuMV activates and manipulates NBR1-ATG8f autophagy in an UPR-dependent manner to anchor VRC to the tonoplast to promote viral replication and virion accumulation (45). Interestingly, TuMV P1 protein interacts with a chloroplast protein cpSRP54 to mediate its degradation via the ubiquitin-proteasome and autophagy pathways to suppress jasmonic acid (JA) biosynthesis and enhance viral infection (46). Therefore, the complicated interactions between potyvirus and autophagy pathway deserve further investigations.

As summarized, potyviral 6K1 is an understudied viral unit, in particular that its expression profile and biological relevance await further investigations. We performed a comprehensive alanine substitution screening, and identified 15 conserved residues in 6K1 sequence that are essential for the infection of pepper veinal mottle virus (PVMV, *Potyvirus* genus). However, 6K1 protein undergoes a degradation event and is less accumulated in virus-infected cells. We demonstrated that cellular autophagy is engaged in 6K1’s degradation. Moreover, we identified four residues enclosed in a conserved lysine/arginine-rich motif in potyviral 6K1s, which are engaged in the autophagy-mediated degradation for the promotion of viral infection.

## RESULTS

### Construction of a GFP-tagged PVMV clone

For potyviruses, both P1/HC-Pro and NIb/CP intercistronic sites are widely engineered to express heterologous proteins (4, 47). To visually monitor PVMV infection, we employed the infectious cDNA clone of the isolate PVMV-HNu (termed pHNu) (48) as the backbone to integrate a complete *GFP* sequence into NIb/CP junction. The resulting clone was designated as pHNu-GFP (Fig. 1A). The original cleavage site ‘DFVLHQ/AG’ at NIb/CP junction (recognized by NIa-Pro) was introduced into both NIb/GFP and GFP/CP junctions for the release of free GFP along with viral genome expression. To examine the infectivity of pHNu-GFP, *N. benthamiana* and *Capsicum chinense* seedlings (*n* = 8 per plant species) were inoculated with pHNu-GFP via agro-infiltration. At 5 days post-inoculation (dpi), *N. benthamiana* plants started to show green fluorescence signals along veins in top non-inoculated leaves under UV lamp (Fig. 1B). The strong fluorescence signals were observed in top leaves for all inoculated plants at 10 dpi and 30 dpi (Fig. 1B). For all *C. chinense* plants inoculated with pHNu-GFP, obvious fluorescence signals were shown at 10 dpi and 15 dpi (Fig. 1C). Similar with pHNu (48), pHNu-GFP induces severe symptoms (such as foliar chlorosis and rugosity, and dwarfism in size) in both *N. benthamiana* and *C. chinense*. Furthermore, the upper non-inoculated leaves of diseased *N. benthamiana* and *C. chinense* plants were harvested for immunodetection of GFP. As anticipated, a major band corresponding to the putative size of free GFP (∼27.7 kDa) was detected in infected leaves (Fig. 1D), indicating that free GFP is efficiently processed and released from viral genome-encoded polyprotein by NIa-Pro.

**Fig. 1.**
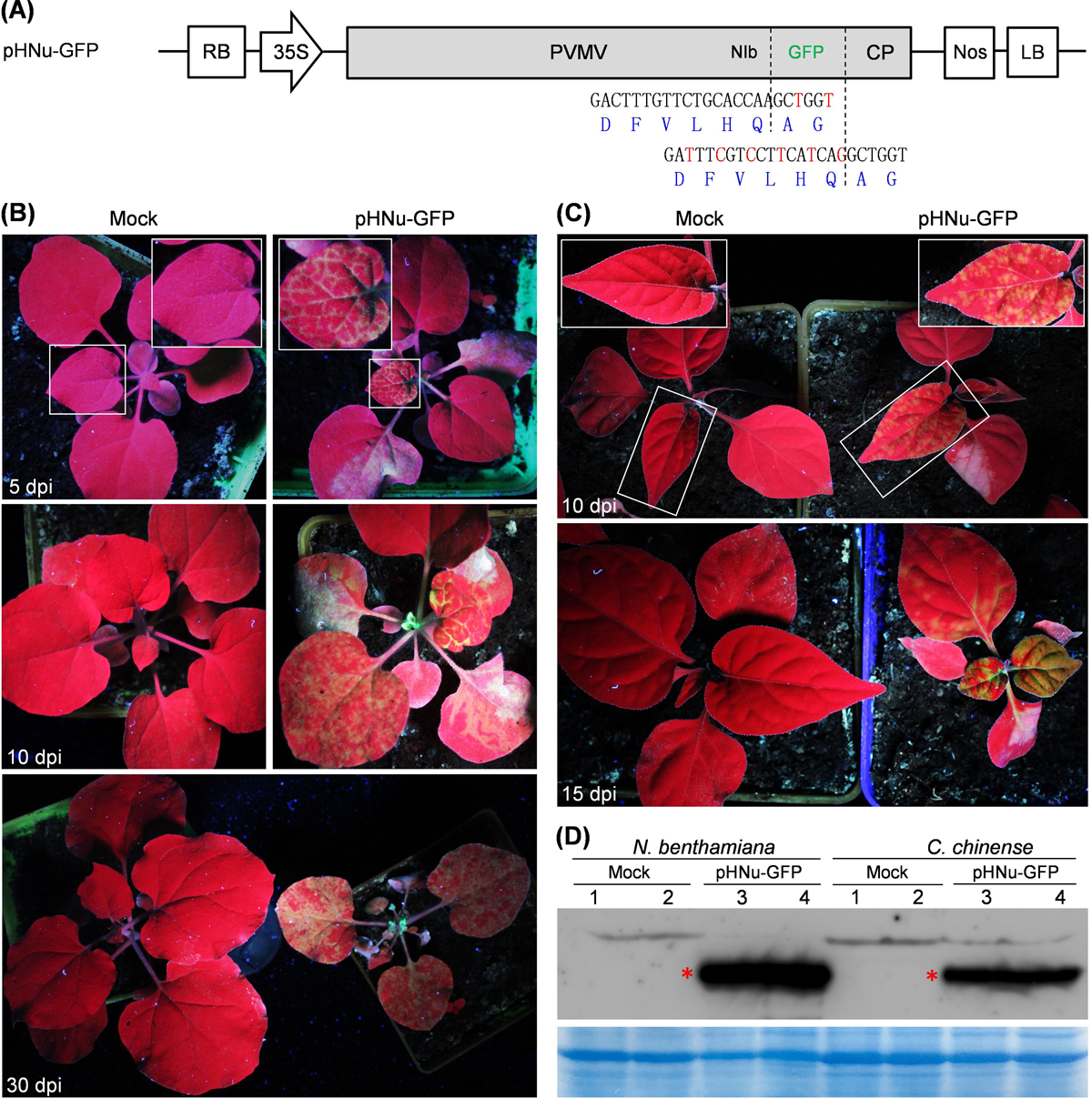
Infectivity test of pHNu-GFP in *N. benthamiana* and *C. chinense*. (A) A schematic diagram of GFP-tagged PVMV clone (pHNu-GFP). A complete GFP-coding sequence was engineered into NIb/CP junction of pHNu (48) to produce the recombinant clone pHNu-GFP. For pHNu-GFP, the original cleavage site ‘DFVLHQ/AG’ at NIb/CP junction was respectively integrated into NIb/GFP and GFP/CP junctions. The mutated nucleotides (in red) without altering cleavage peptide sequence were introduced to avoid the removal of *GFP* sequence via recombination event during viral replication. (B) Infectivity test of pHNu-GFP in *N. benthamiana*. The representative plants were photographed under UV lamp. Mock, empty vector control, pCB301. The close-view of indicated regions by rectangles is shown. (C) Infectivity test of pHNu-GFP in *C. chinense*. (D) Immunoblotting detection of free GFP in top non-inoculated leaves of *N. benthamiana* and *C. chinense* plants. The bands indicated by asterisks correspond to the predicted size of free GFP (∼27.7 kDa). A Coomassie brilliant blue-stained gel was used as a loading control. Two samples per treatment were assayed.

### Fifteen out of 17 conserved residues across potyviral 6K1s are essential for PVMV infection in *N. benthamiana* or *C. chinense*

To investigate the biological significance of 6K1 sequence during viral infection, a total of 115 sequences of 6K1 from different potyviruses were retrieved from NCBI GenBank database, and subjected to multiple alignment analysis. The results showed that potyviral 6K1 sequences are rather conserved. A total of 17 highly-conserved residues, excluding the conserved Gln at the position P1 of NIa-Pro cleavage site at 6K1-CI junction, were characterized: Lys / Arg (K3, 38; K/R28, 34, 40), Asp / Glu (E11; D25, 30; D/E27), Ala (A15), Leu (L19, 36, 39), Met (M22), Ser (S29), and Val (V32, 51) (Fig. 2A). We performed alanine substitution screening to evaluate the effects of these conserved residues on viral infectivity. For A15, it was substituted with Arg. Using pHNu-GFP as the backbone, we produced a total of 17 mutated clones, by which the resulting virus mutants are expected to harbor single substitution of conserved residues in 6K1 with Ala, or A15 with Arg. For the convenience of description, pHNu-GFP is designated as WT in this section. These mutated clones, together with WT, were individually inoculated into both *N. benthamiana* (*n* = 8 per clone) and *C*. *chinense* seedlings (*n* = 5 per clone) by agro-infiltration.

**Fig. 2.**
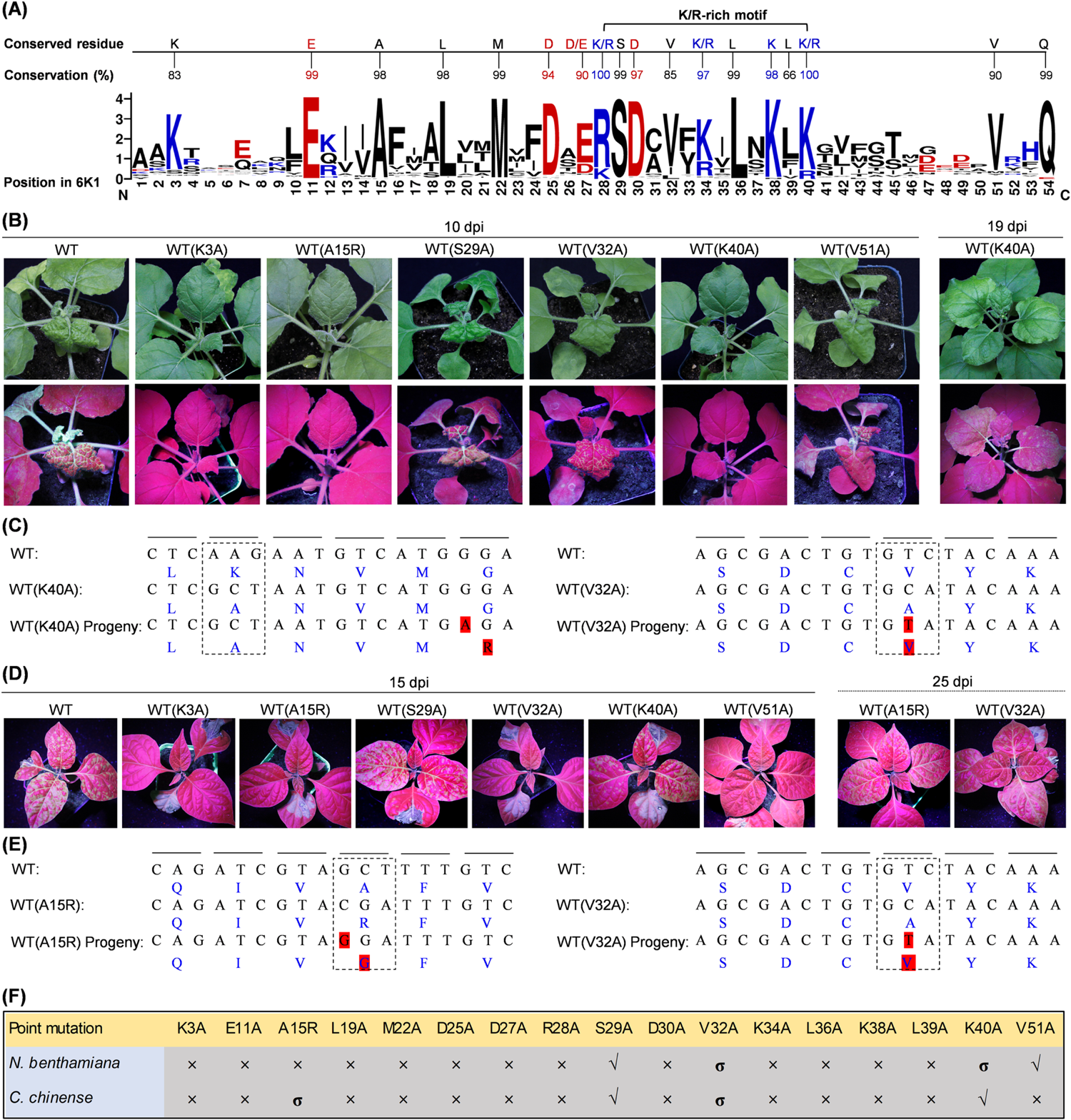
The effects of 17 different point substitutions in PVMV 6K1 on viral infectivity in both *N. benthamiana* and *C. chinense*. (A) The analysis of amino acid conservation of 115 sequences of 6K1 from different potyviruses. The order of amino acids was sorted with reference to PVMV 6K1. Alkaline amino acids are shown in blue, acidic ones in red, and the rest in black. (B, D) Infectivity test of mutated PVMV clones in *N. benthamiana* (B) and *C. chinense* (D). A close-view of representative plants is shown. The GFP signals were examined under an handheld UV lamp in a darkness room at the indicated time points. (C, E) Sequencing analysis of virus progeny. Sequence comparison across WT, mutated clones, and viral progeny. The mutated sites and surrounding sequences are shown. The sequences of viral progeny derived from WT(V32A) and WT(K40A) in *N. benthamiana* was determined at 19 dpi (C). The sequences of viral progeny derived from WT(V32A) and WT(A15R) in *C. chinense* were determined at 25 dpi. The amino acids corresponding to the codons are shown in blue, and reversion and compensatory mutations are shaded in red. (F) Summary on the infectivity of different mutated clones in *N. benthamiana* and *C. chinense*. √, the mutated clones that are able to efficiently infect plants; ×, the clones that are disabled in successful infection; σ, the clones that are attenuated in systemic infection.

In *N. benthamiana*, all plants inoculated with WT(S29A) or WT(V51A) exhibited severe distortion symptoms and strong GFP signals in top leaves at 10 dpi, resembling those inoculated with WT. Two of five plants inoculated with WT(V32A) showed mild symptoms and weak GFP signals (Fig. 2B). Neither virus-infected symptoms nor GFP signals were observed on plants inoculated with the other mutated clones at this time point (Fig. 2B, 2F, and Fig. S1). Intriguingly, clear GFP signals started to show in top leaves of three out of five plants inoculated with WT(K40A) at 19 dpi (Fig. 2B). Except the plants mentioned above, the remaining ones did not show any discernible symptom or GFP signals, even until 30 dpi. For virus progeny derived from WT(S29A), WT(V32A), WT(K40A), and WT(V51A), the genomic sequence, covering P3 C-terminus, 6K1, and CI N-terminus, was determined. Spontaneous mutations were not found for both WT(S29A) and WT(V51A). However, one reversion mutation ‘C to T’, leading to ‘A to V’ at position 32 was identified for the progeny of WT(V32A) (Fig. 2C). For the progeny of WT(K40A), a compensatory mutation ‘G to A’, resulting in ‘G to R’ at position 44, took place (Fig. 2C). Taken together, the above results confirm that all conserved residues across potyviral 6K1s, excluding S29 and V51, are key for a successful infection of PVMV in *N. benthamiana*.

In *C. chinense*, all plants inoculated with WT(S29A) or WT(K40A) displayed strong GFP signals in top leaves at 15 dpi, resembling WT-inoculated plants (Fig. 2D). Intriguingly, strong GFP signals appeared in top leaves of two WT(V32A)-inoculated and one WT(A15R)-inoculated plants at 25 dpi (Fig. 2D). Except the plants mentioned above, the remaining plants did not show GFP signals (Fig. 2D, 2F, and Fig. S2), even until 45 dpi. Similarly, the genomic sequences surrounding 6K1 for viral progeny were determined. No spontaneous mutations were found for the progeny of WT(S29A) and WT(K40A). In line with the observation in *N. benthamiana*, the reversion mutation ‘A to V’ at position 32 was found for the progeny of WT(V32A) (Fig. 2E). For the progeny of WT(A15R), one mutation ‘C to G’, resulting in ‘R to G’ at position 15, was observed (Fig. 2E). Collectively, all conserved residues in 6K1s, with the exception of S29 and K40, are essential for PVMV infection in its natural host, *C. chinense*.

### 6K1 protein is less accumulated during PVMV infection

To examine the expression dynamics of mature 6K1 in viral infection, we created a modified virus clone, pHNu-GFP-6K1^Myc^, to express a C-terminal Myc-fused 6K1 (Fig. 3A). Infectivity test showed that the plants inoculated with pHNu-GFP-6K1^Myc^ exhibited similar symptoms and distribution pattern of GFP signals with those treated with pHNu-GFP in *N. benthamiana* (Fig. 3B). The presence of virus in upper non-inoculated leaves was confirmed by RT-PCR (Fig. 3C). For the virus progeny, the sequence spanning the Myc tag was determined, and spontaneous mutations / deletions were not observed (Fig. 3D), indicating that the attachment of Myc epitope at the C-terminus of 6K1 has no obvious effect on viral infectivity. Both inoculated (IL) and upper non-inoculated leaf (TL) samples were harvested at different time points, and subjected to immunodetection of 6K1^Myc^. The results showed that a specific band corresponding to the putative size of the precursor P3-6K1^Myc^ (∼47.0 kDa) was detected not only from IL samples at 5 dpi and 6 dpi, but also from TL samples at 5 dpi, 6 dpi, and 7 dpi (Fig. 3E). However, the mature 6K1^Myc^ (∼7.6 kDa) was not immuno-detected at all these time points (Fig. 3E), implying that a low abundance of 6K1 protein, at an undetectable level by immunoblotting, is along with viral infection.

**Fig. 3.**
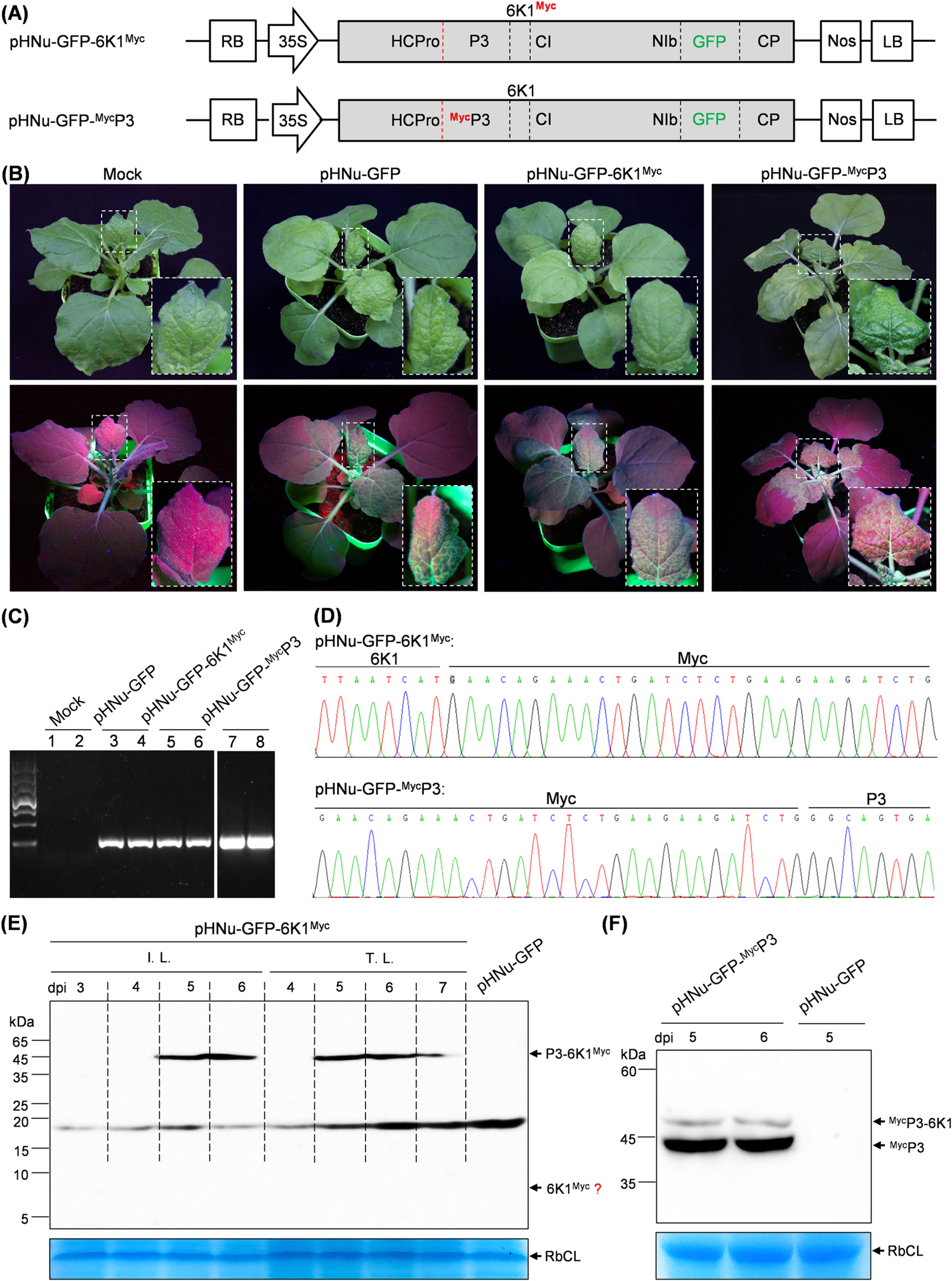
The expression profile of 6K1 during PVMV infection. (A) Schematic diagrams of pHNu-GFP-^Myc^P3 and pHNu-GFP-6K1^Myc^. The dotted line in red denotes the self-cleavage site of HCPro, and the white ones represent the cleavage sites by NIa-Pro. (B) Infectivity test of pHNu-GFP-^Myc^P3 and pHNu-GFP-6K1^Myc^. The GFP signals in inoculated *N. benthamiana* plants are examined under a handheld UV lamp at 8 dpi. The close-up of leaves in white rectangle are shown. Mock, empty vector control. (C) RT-PCR detection of viral infection in inoculated plants. The upper non-inoculated leaves were sampled at 8 dpi for RT-PCR detection with primer set PVMV-F/PVMV-R (48) targeting viral *CP* cistron. (D) Sequencing analysis of virus progeny derived from pHNu-GFP-6K1^Myc^ and pHNu-GFP-^Myc^P3. The upper non-inoculated leaves were harvested at 8 dpi, and used for cloning and sequencing. (E) Immunoblot detection of the expression profile of 6K1 in viral infection. I.L., inoculated leaves; T.L., upper non-inoculated leaves. Immunoblot analysis of the sample from upper non-inoculated leaves of pHNu-GFP at 8 dpi was included as the negative control. A Coomassie brilliant blue-stained gel was used as a loading control. (F) Immunoblotting analysis of the expression of ^Myc^P3 and ^Myc^P3-6K1 in viral infection. The upper non-inoculated leaves were sampled for the assay at the indicated time points. Immunoblot analysis of the sample from upper non-inoculated leaves of pHNu-GFP at 8 dpi was included as the negative control. A Coomassie brilliant blue-stained of Rubiso large unit (RbCL) was used as a loading control.

### P3-6K1 junction is efficiently processed by NIa-Pro in viral infection

Previously studies based on *in vitro* assays or in insect cells showed that 6K1-CI junction in potyviral polyprotein was efficiently processed by NIa-Pro but the processing at P3-6K1 junction at a low rate (25–27). These observations compeled us to speculate that a low abundance of 6K1 during PVMV infection might be the consequence of low-efficient cleavage by NIa-Pro at P3-6K1 junction. To test this idea, we examined the processing efficiency of P3-6K1 junction in viral infection. We created a modified virus clone, pHNu-GFP-^Myc^P3, to express N-terminal Myc-fused P3 or P3-6K1 (if have) (Fig. 3A). The clone was inoculated into *N. benthamiana* seedlings (*n* = 10). The results showed that pHNu-GFP-^Myc^P3 behaved like pHNu-GFP, in terms of symptom phenotype and GFP distribution pattern in upper non-inoculated leaves (Fig. 3B). Viral infection in upper non-inoculated leaves were confirmed by RT-PCR (Fig. 3C). The sequence spanning the Myc tag for virus progeny was determined, and spontaneous mutations / deletions were not found (Fig. 3D), indicating that the fusion of Myc epitope with the N-terminus of P3 has no discernible effect on viral infectivity. Leaf samples were harvested from upper non-inoculated leaves at 5 and 6 dpi, and used for immunodetection with anti-Myc polyclonal antibody. Two specific protein bands with the putative size for ^Myc^P3-6K1 (∼46.8 kDa) and ^Myc^P3 (∼40.7 kDa) were detected (Fig. 3F). However, the signal intensity of the band representing ^Myc^P3 was markedly higher than that of ^Myc^P3-6K1 (Fig. 3F), indicating that P3-6K1 junction is efficiently processed by NIa-Pro during PVMV infection. In combination of the data in previous section, it is suggesting that PVMV 6K1 is efficiently released from viral polyprotein, but undergoes an intracellular self-degradation event.

### Proteolytic processing at P3-6K1 junction is indispensable for the successful infection of PVMV

Next, we tested whether the processing of viral polyprotein at P3-6K1 junction is required for viral infection. Previous studies based on *in vitro* assays showed that amino acid substitution in conserved heptapeptide recognized by NIa-Pro alters or disables the efficiency of proteolytic processing (26, 49). Substitution of Gln with His at P1 position blocks the processing by NIa-Pro (26, 50). Replacement with Lys at P1′ position disturbs NIa-Pro processing (51). Statistically, Gln is the most frequent residue at P1 position, whereas Ala is absent at this position. At P1′ position, Ala, Ser and Gly are the most frequent residues, but Gln is absent (49).

Based on the above observations, five substitutions, Q-H, A-K, QA-AQ, Q-A, and A-Q, were individually introduced into the heptapeptide at P3-6K1 junction by using pHNu-GFP-^Myc^P3 as the backbone, to destroy the cleavage of NIa-Pro at P3-6K1 junction (Fig. 4A). All mutated clones were inoculated into *N. benthamiana* seedlings (*n* = 8 per clone). As shown in Fig. 4B, sorely pHNu-GFP-^Myc^P3(A-Q), behaving like PVMV-GFP-^Myc^P3, was aggressive in *N. benthamiana*. Further, we examined the cleavage of NIa-Pro at P3-6K1 junction for these mutants *in planta*. The inoculated leaves were harvested at 5 dpi for immunoblotting detection with anti-Myc polyclonal antibody. The results showed that both ^Myc^P3 and ^Myc^P3-6K1 were detected for pHNu-GFP-^Myc^P3(A-Q) and pHNu-GFP-^Myc^P3 (Fig. 4D), indicating that the P3-6K1 junction for them is processed by NIa-Pro in viral infection. Noticeably, the signal intensity of the band corresponding to ^Myc^P3 is much weaker than that of P3-6K1 for PVMV-GFP-^Myc^P3(A-Q), suggesting a relatively low processing efficiency at the mutated cleavage site. Unfortunately, either P3-6K1 or P3 was not detected for the other four mutants (Fig. 4D). A most likely explanation is that these introduced mutations prevent NIa-Pro’s cleavage at P3-6K1 junction, compromise viral multiplication, and contribute to the accumulation of P3-6K1 at an undetectable level.

**Fig. 4.**
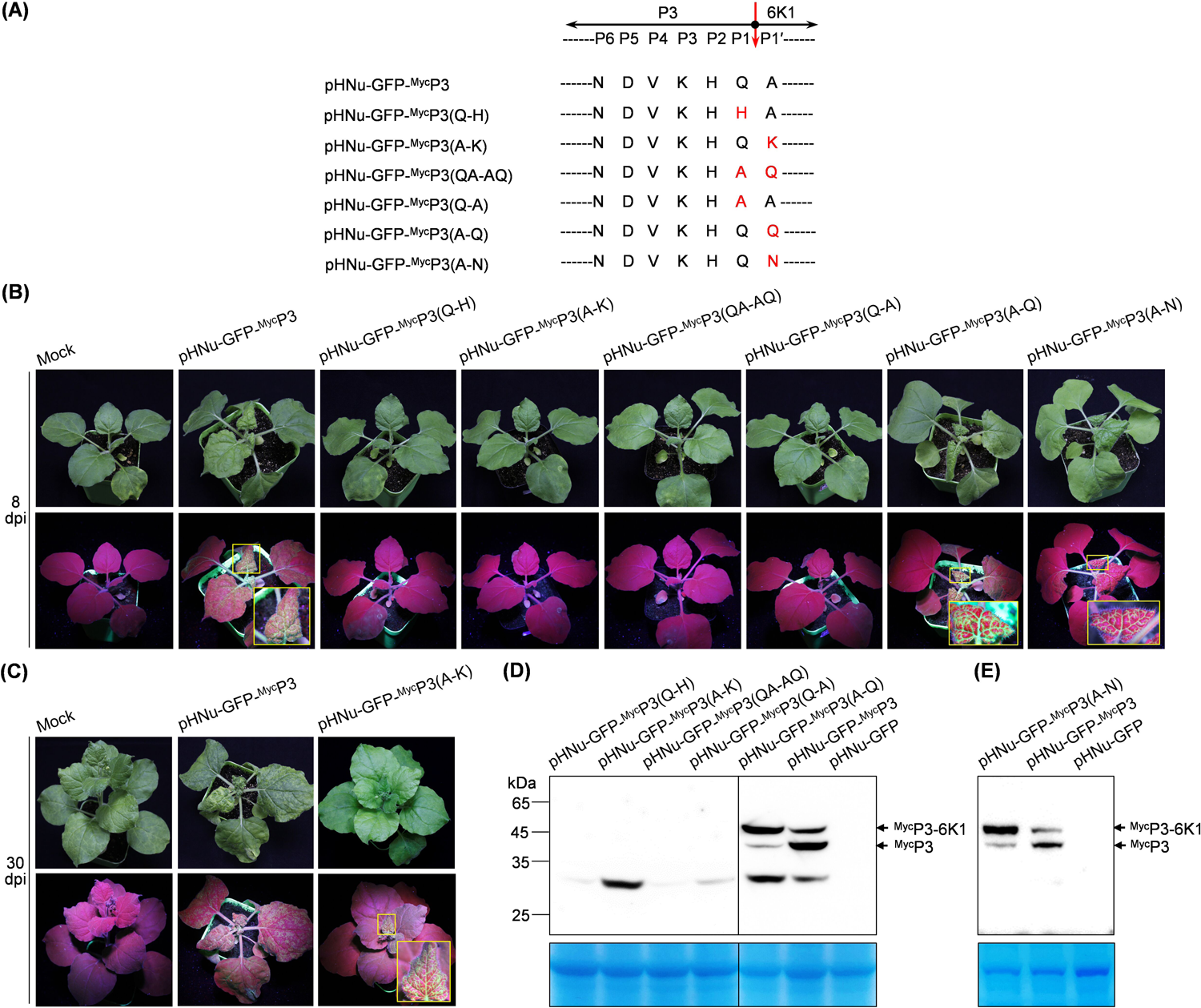
Proteolytic processing at P3-6K1 junction by NIa-Pro is indispensable for the successful infection of PVMV. (A) Schematic diagram showing the mutations introduced into conserved heptapeptide at P3-6K1 junction. NIa-Pro cleavage site between P3 and 6K1 is indicated by a red arrow. The amino acids of heptapeptide were positioned as P1 through P6 and P1′, with reference to a previous document (49). The introduced mutated amino acids are shown in red. (B, C) The effects of different mutations at the cleavage site between P3 and 6K1 on viral infectivity in *N. benthamiana*. Representative plants were photographed at 8 dpi (B) or 30 dpi (C). The GFP signals in each representative plant were examined under a handheld UV lamp in a darkness room. The leaf regions in yellow rectangle are enlarged. Mock, empty vector control. (D, E) Immunoblotting analysis of the expression of P3 and P3-6K1. Leaf patches inoculated with the indicated clones were sampled at 5 dpi, and were used for immunoblot detection of P3 and P3-6K1 by using an anti-Myc polyclonal antibody. A Coomassie brilliant blue-stained gel was used as a loading control.

Intriguingly, GFP signals started to emerge in top leaves of one plant inoculated with pHNu-GFP-^Myc^P3(A-K) at 21 dpi, and became strong at 30 dpi (Fig. 4C), indicating that virus progeny derived from this clone are infectious in *N. benthamiana*. For the virus progeny, the sequence surrounding P3-6K1 junction was determined. A spontaneous nucleotide mutation, leading to the reversion from Lys to Asn at P1′ position, was detected. We proposed that the spontaneous mutation might recover the cleavage of NIa-Pro at P3-6K1 junction, leading to viral successful infection. To test this idea, the substitution with Asn at P1′ position was introduced (Fig. 4A) to generate the clone pHNu-GFP-^Myc^P3(A-N). The infectivity of the clone was greatly recovered, evidenced by that all eight inoculated plants exhibited severe leaf rugosity symptoms and strong GFP signals in top leaves, resembling the plants inoculated with pHNu-GFP-^Myc^P3 (Fig. 4B). The expression of both P3 and P3-6K1 was examined. The result showed that both proteins were immuno-detected, indicating the reversion mutation ‘K to N’ at P1′ position recover, albeit partially, the cleavage at P3-6K1 junction, and, thus, rescue viral infectivity.

Collectively, the above results support the notion that proteolytic processing at P3-6K1 junction is indispensable for viral successful infection. In combination with the observation that 6K1 is less accumulated in viral infection (Fig. 3), it is speculated that the generation of mature 6K1 along with its intracellular degradation might be important for viral infection.

### PVMV 6K1 undergoes autophagic degradation

Potyviral 6K1s, when ectopically expressed *in planta*, diffuse into cytoplasm and nucleus (30, 31, 52, 53). Once expressed in the context of viral infection, they would form functional punctate structures targeting 6K2-induced replication vesicles (30, 31). This prompted us to investigate, in the context of viral infection, the mechanism by which PVMV 6K1 is degraded. For the convenience of the detection of 6K1, we engineered a second copy of 6K1 with its C-end fused with GFP into NIb-CP junction of pHNu to generate a recombinant clone, pHNu//6K1-GFP) (Fig. 5A). The attachment of a GFP-tag to the C-terminus of 6K1 seems not affect its colocalization with 6K2-induced replication vesicles (30). The recombinant clone enables our detection of 6K1 in the context of viral infection. Autophagy and ubiquitin-proteasome machineries are two main intracellular degradation pathways (54, 55). We selected two autophagy inhibitors, 3-methyladenine (3-MA) and E-64d, and a 26S proteasome inhibitor MG132 to determine the effects of autophagy and ubiquitin-proteasome pathways on the degradation of 6K1. Agrobacterial cultures harboring pHNu-GFP (as the parallel control) or pHNu//6K1-GFP (OD_600_ = 1.0 per culture) were primarily inoculated into fully-expanded leaves of *N. benthamiana* seedlings at 6- to 8-leaf stage. At 80 hours post-inoculation (hpi), the inoculated leaves were treated with different chemical inhibitors (Fig. 5B). Sixteen hours later, the treated leaves were sampled for immunoblotting analysis. E-64d treatment significantly increased the accumulation of 6K1-GFP, whereas the treatment did not obviously alter the accumulation of free GFP in the parallel control (Fig. 5C). The results suggest that cellular autophagy is engaged in the degradation of 6K1 in viral infection. Even though MG132 treatment enhanced the abundance of 6K1-GFP, the enhancing effect was also observed for free GFP in the parallel control (Fig. 5C). Herein, it could not be concluded whether the ubiquitin-proteasome pathway is engaged in 6K1’s degradation. In contrast with E-64d, the treatment with 3-MA had no increasement effect on the accumulation of 6K1-GFP (Fig. 5C).

**Fig. 5.**
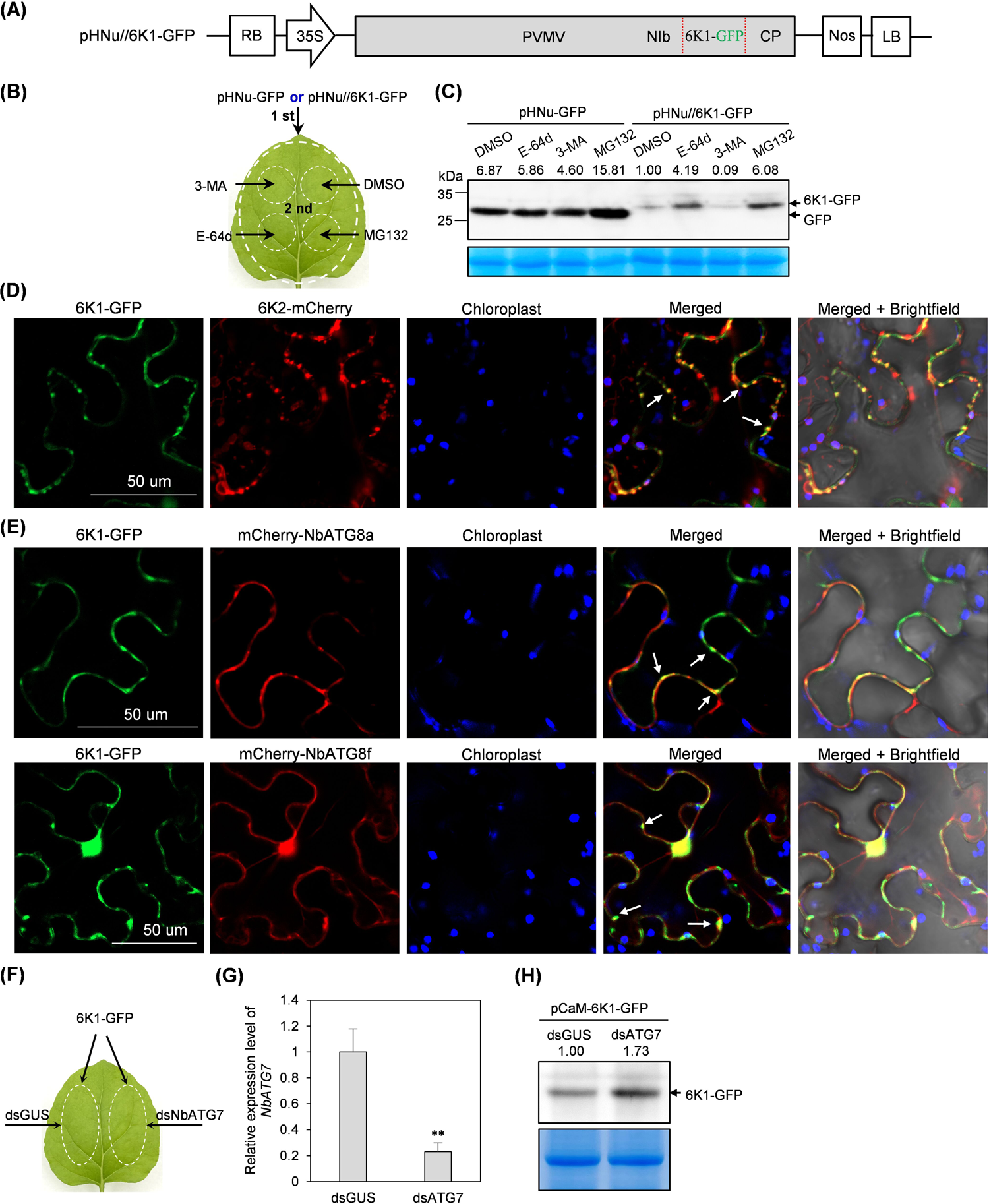
PVMV 6K1 is degraded by cellular autophagy. (A) Schematic diagram of pHNu//6K1-GFP. The red lines flanking 6K1-GFP represent NIa-Pro’s cleavage sites at NIb-6K1-GFP and 6K1-GFP-CP junctions. (B) The patch design for the combinations of agrobacterial infiltration and chemical inhibitor treatment in the same leaves. 1 st, primary infiltration with agrobacterial culture harboring pHNu-GFP or pHNu//6K1-GFP; 2 nd, treatment with chemical inhibitor or DMSO. (C) The effects of ubiquitin-proteasome and autophagy pathways on the degradation of 6K1 in the context of viral infection. The signal intensity of protein bands is shown above the panel. The value for the combination of pHNu//6K1-GFP and p2300s-intron-dsGUS is designated as 1.0 to normalize the data. A Coomassie brilliant blue-stained gel was used as a loading control. (D) Co-localization of 6K1-GFP and 6K2-mCherry in the context of viral infection. Arrows, overlapped structures of 6K1-GFP and 6K2-mCherry. Bar, 50 μm. (E) Co-localization of 6K1-GFP and mCherry-NbATG8a or mCherry-NbATG8f at 60 hpi. Arrows, overlapped structures of 6K1-GFP and mCherry-NbATG8a / mCherry-NbATG8f. Bars, 50 μm. (F) The patch design for co-infiltration with agrobacterial cultures harboring pCaM-6K1-GFP and p2300s-intron-dsATG7 / p2300s-intron-dsGUS. (G) RT-qPCR analysis of the abundance of *NbATG7* mRNA transcripts at 3 dpi. The expression level of *NbActin* transcripts was determined to normalize the data. Error bars denote standard errors from three biological replicates. Statistically significant differences, determined by an unpaired two-tailed Student’s *t* test, are indicated by asterisks: **, 0.001<*P*< 0.01. (H) Effect of *NbATG7* knockdown on the accumulation of 6K1-GFP. Western blot analysis of the abundance of 6K1-GFP at 3 dpi. A Coomassie brilliant blue-stained gel was used as a loading control.

Previous studies reported that potyviral 6K1s form punctate structures that target 6K2-induced VRC in virus-infected cells (30, 31). The colocalization of PVMV 6K1 with 6K2-induced VRC was tested. Agrobacterial culture harboring pHNu//6K1-GFP (0.5 of OD_600_) was infiltrated into *N. benthamiana* leaves, which, at 24 hpi, were re-inoculated with an agrobacterial culture harboring pCaM-6K2-mCherry (0.3 of OD_600_). Thirty-six hours later, 6K1-GFP forms punctate structures, which largely co-localize with 6K2-mCherry vesicles (Fig. 5D). When autophagy pathway is activated, ATG8 proteins ectopically expressed *in planta* would form punctate structures representing autophagosomes (42, 56, 57). Both ATG8a and ATG8f from *N. benthamiana* with their N-ends fused with mCherry (mCherry-NbATG8a and mCherry-NbATG8f) were used as autophagosome markers to further dissect the association of 6K1 with autophagic event. The agrobacterial culture harboring pHNu//6K1-GFP, together with the culture containing pCaM-mCherry-NbATG8a or pCaM-mCherry-NbATG8f (final OD_600_ = 0.25 per culture), were co-inoculated into fully-expanded leaves of *N. benthamiana* seedlings at 6- to 8-leaf stage. Indeed, a portion of punctate structures formed by 6K1-GFP co-localized with mCherry-NbATG8a or mCherry-NbATG8f structures in cytoplasm (Fig. 5E). These data indicate that 6K1 is physically associated with ATG8s-labled autophagosomes during viral infection.

To further determine the degradation of 6K1 by cellular autophagy, we constructed one hairpin RNAi construct (p2300s-intron-dsATG7) for the transient silencing of *NbATG7* to block cellular autophagy pathway, and another one p2300s-intron-dsGUS as the parallel control. Meanwhile, a T-DNA construct (pCaM-6K1-GFP) for transient expression of 6K1-GFP was developed. Agrobacterial cultures, harboring corresponding pCaM-6K1-GFP and p2300s-intron-dsATG7 or p2300s-intron-dsGUS (final OD_600_ = 0.3 per culture) were co-infiltrated into fully-expanded leaves of *N. benthamiana* seedlings at 6- to 8-leaf stage (Fig. 5F). Seventy-two hours later, the mRNA transcripts of *NbATG7* in inoculated leaves with the combination of p2300s-intron-dsATG7 and pCaM-6K1-GFP was reduced by 76%, compared with the parallel control (Fig. 5G). In turn, an increasing amount of 6K1-GFP by 73%, in comparison with that from the parallel control, was detected (Fig. 5H).

### Individual expression of five 6K1 variants (D30A, V32A, K34A, L36A, and L39A) along with viral infection significantly interfere with viral infection progression; the five residues are encompassed in a Lys/Arg-rich motif across potyviral 6K1s

Herein, we hypothesized that: ⅰ) the degradation of 6K1 by cellular autophagy might be important for viral infection; ⅱ) there might be the existence of potential amino acid(s) or motif(s) in 6K1 that determine its autophagic degradation, and mutating these residues would interfere with viral infectivity. To test the ideas, the fifteen *6K1* variants (Fig. 2) were individually engineered into pHNu at the intercistronic junction of *NIb* and *CP* (Fig. 6A). Each of the resulting recombinant clones were inoculated into three *N. benthamiana* seedlings at 9- to 10-leaf stage (OD_600_ = 0.1 per clone). At 5 dpi, all plants inoculated with pHNu//6K1 exhibited obvious vein-clearing and distortion symptoms in top leaves (Fig. 6B). Similar symptoms were also observed in plants inoculated with pHNu//6K1 variants (K3A, E11A, A15R, M22A, D25A, D27A, and K38A) (Fig. S3A). However, virus-induced symptoms were not discernible in plants inoculated with pHNu//6K1 variants (L19A, R28A, D30A, V32A, K34A, L36A, L39A, and K40A) at this time point (Fig. 6B). In line with above observations, a significantly lower abundance of viral genomic RNAs was detected from plants inoculated with these clones, with the exception of pHNu//6K1(L19A) (Fig. 6C). It was noticed that all pHNu//6K1 variants, behaving like pHNu//6K1, induced nearly-consistent severe symptoms at 7 dpi (Fig. S3B). Consistently, a comparable level of viral RNA accumulation was shared by all pHNu//6K1 variants and pHNu//6K1 at this time point (Fig. S3C). Conclusively, individual expression of seven 6K1 variants (R28A, D30A, V32A, K34A, L36A, L39A, and K40A) significantly delays viral infection progression in *N. benthamiana*, likely via interfering with the function of the cognate 6K1 in PVMV. Further, the performance of the seven recombinant clones in *C. chinense* was investigated. These pHNu//6K1 variants, together with pHNu//6K1, were each inoculated into six *C. chinense* plants at 5- to 6-leaf stage (OD_600_ = 0.1 per clone). All plants inoculated with pHNu//6K1(R28A), or pHNu//6K1(K40A), similar with the plants inoculated with pHNu//6K1, exhibited obvious virus-induced symptoms at 10 dpi, such as chlorosis along veins in upper non-inoculated leaves (Fig. 6D). In contrast, no or mild symptoms were observed in plants inoculated with pHNu//6K1 variants (D30A, V32A, K34A, L36A, and L39A) (Fig. 6D). In accordance, a significantly lower viral RNA accumulation level was detected for the groups of plants showing no or mild symptom (Fig. 6E). Similar with what was observed in *N. benthamiana*, all *C. chinense* plants inoculated with each of pHNu//6K1variants or pHNu//6K1 exhibited indistinguishable severe symptoms at 15 dpi (Fig. S3D). A comparable level of viral RNA accumulation was shared by these pHNu//6K1 variants and pHNu//6K1 at this time point (Fig. S3E).

**Fig. 6.**
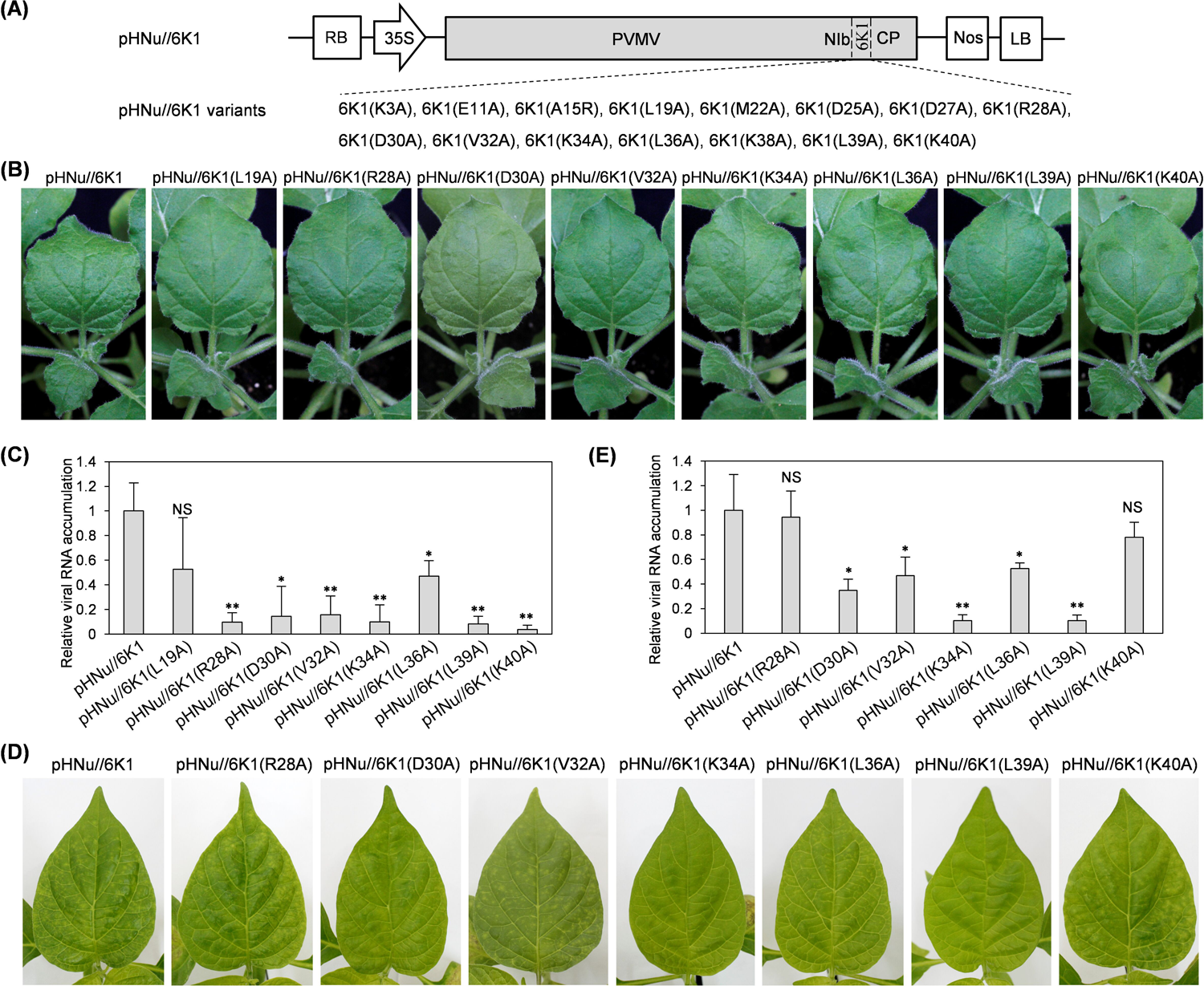
Individual expression of five 6K1 mutants along with viral infection delays the infection progression in both *N. benthamiana* and *C. chinense*. (A) Schematic diagram of pHNu//6K1 and pHNu//6K1 variants. (B, D) Infectivity test of pHNu//6K1 variants in both *N. benthamiana* and *C. chinense*. Photographs were taken at 5 dpi for *N. benthamiana* (B), and at 10 dpi for *C. chinense* (D). (C, E) RT-qPCR analysis of viral accumulation levels. The upper non-inoculated leaves of *N. benthamiana* at 5 dpi (C) and *C. chinense* at 10 dpi (E), were sampled for RT-qPCR assays. The expression levels of *NbActin* and its ortholog in *C. chinense* (*CcActin*) transcripts were determined to normalize the data. Error bars denote standard errors from three biological replicates. Statistically significant differences, determined by an unpaired two-tailed Student’s *t* test, are indicated by asterisk: **, 0.001<*P*< 0.01; *, 0.01<*P*< 0.05.

Taken together, individual expression of five 6K1 variants (D30A, V32A, K34A, L36A, and L39A) along with viral infection greatly hinders viral infection progression in both *N. benthamiana* and *C. chinense*. Further analysis revealed that the five conserved residues are encompassed in a Lys/Arg-rich motif across potyviral 6K1s (Fig. 2A).

### The four conserved residues (V32, K34, L36, L39) across potyviral 6K1s are related with the autophagic degradation of PVMV 6K1

Next, we investigated whether the five conserved residues (D30, V32, K34, L36, and L39) are engaged in the autophagic degradation of PVMV 6K1. To test this hypothesis in the context of viral infection, individual substitution of them with alanine was introduced into 6K1-GFP in pHNu//6K1-GFP (Fig. 7A). The obtained clones pHNu//6K1 variants-GFP together with pHNu//6K1-GFP were each inoculated into fully-expanded leaves of *N. benthamiana* leaves at 6- to 8-leaf stage (OD_600_ = 1.0 per culture). At 80 hpi, the inoculated leaves were treated with E-64d or DMSO (Fig. 7B). Sixteen hours later, the treated leaves were sampled for immunoblotting analysis. The results showed that E-64d treatment failed to enhance the accumulation of 6K1(D30A)-GFP, 6K1(V32A)-GFP, 6K1(K34A)-GFP, 6K1(L36A)-GFP, or 6K1(L39A)-GFP, which was in contrast with 6K1-GFP in pHNu//6K1-GFP (Fig. 7C).

**Fig. 7.**
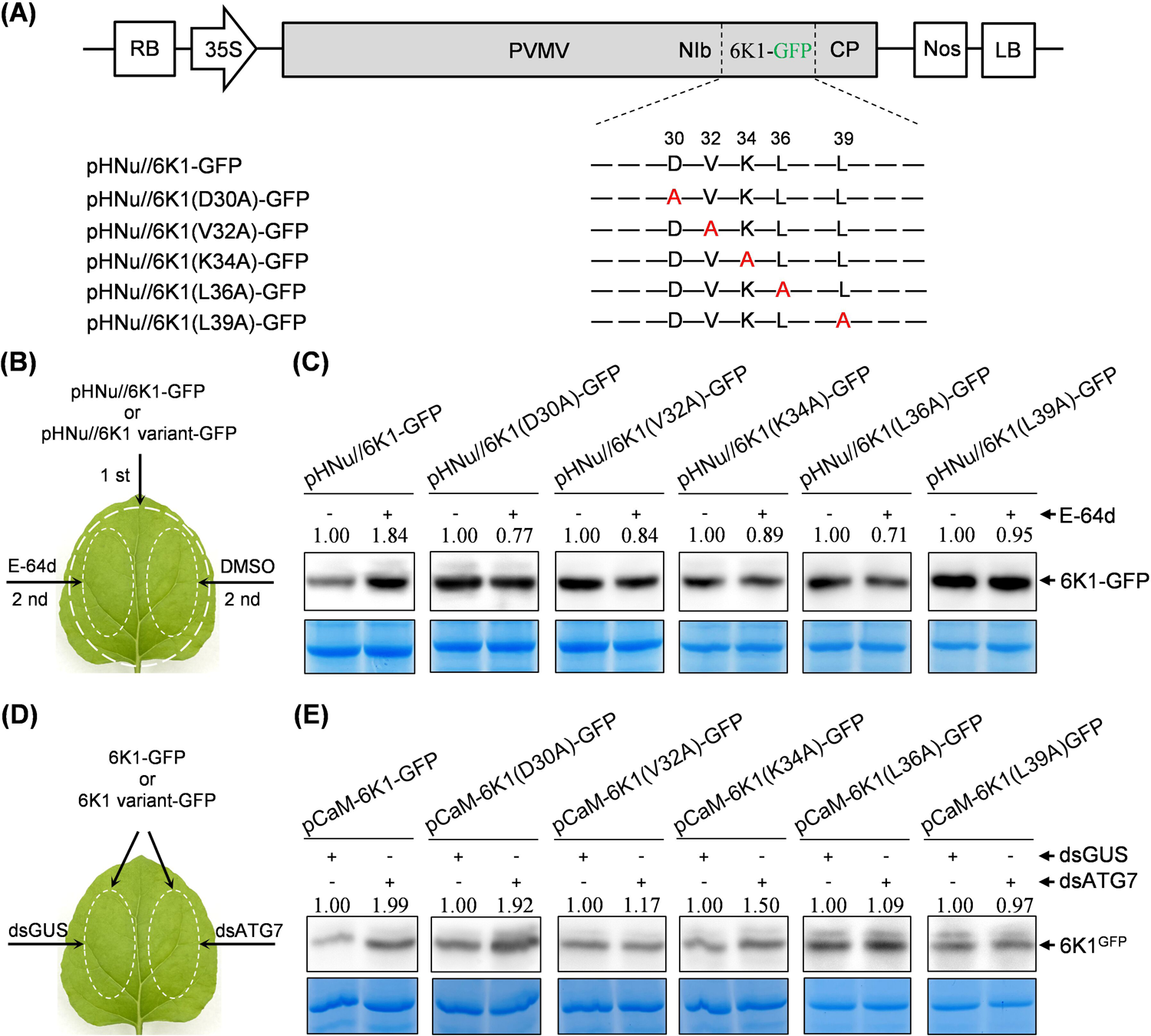
Alanine substitutions of conserved residues (V32, K34, L36, L39) encompassed in a lysine/arginine-rich motif in 6K1 inhibited its autophagic degradation. (A) Schematic diagram of pHNu//6K1 variants-GFP. (B) The patch design for the combinations of agrobacterial infiltration and chemical inhibitor treatment in the same leaves. 1 st, primary infiltration with agrobacterial culture harboring pHNu//6K1-GFP or pHNu//6K1 variants-GFP; 2 nd, treatment with E-64d or DMSO. (C) The effects of E-64d treatment on the accumulation of 6K1 variants in the context of viral infection. The signal intensity of protein bands is shown above the panel. The values for DMSO treatment were designated as 1.0 to normalize the data. -, DMSO treatment; +, E-64d treatment. Coomassie brilliant blue-stained gels were used as loading controls. (D) The patch design for co-infiltration with agrobacterial cultures harboring pCaM-6K1-GFP / pCaM-6K1(point mutaion)-GFP and p2300s-intron-dsATG7 / p2300s-intron-dsGUS. (E) Effect of *NbATG7* knockdown on the accumulation of 6K1-GFP or 6K1 variants-GFP. Western blot analysis of the abundance of 6K1-GFP or 6K1 variants-GFP at 3 dpi. Coomassie brilliant blue-stained gels were used as loading controls.

To further evaluate the association of the five residues with autophagic degradation of 6K1, individual substitution of them with alanine was introduced into 6K1-GFP in pCaM-6K1-GFP. Each agrobacterial culture harboring relevant plasmids was co-infiltrated with p2300s-intron-dsATG7 or p2300s-intron-dsGUS into fully-expanded leaves of *N. benthamiana* seedlings at 6- to 8-leaf stage (Fig. 7D). Seventy-two hours later, the co-inoculated leaves were sampled for immunoblotting analysis. The results showed that the treatment with dsATG7 construct failed to enhance the accumulation of 6K1(V32A)-GFP, 6K1(L36A)-GFP, and 6K1(L39A)-GFP, when compared with the treatment by dsGUS. The dsATG7 treatment increased the amount of 6K1(K34A)-GFP by 50% (Fig. 7E). However, the treatment by dsATG7 construct yielded approximately two-fold amount of 6K1(D30A)-GFP, similar with the case of 6K1-GFP (Fig. 7E). The molecular mechanism underpinning the contrasting effects of D30 on 6K1’s autophagic degradation (transient expression versus the expression in viral infection) (Fig. 7C, 7E), awaits to be further investigated. Collectively, four 6K1 variants (V32A, K34A, L36A, and L39A) inhibited the autophagic degradation of 6K1.

## DISCUSSION

During co-evolution between viruses and plant hosts, the end-less arm races are launched. To survive, viruses continuously forge a limited number of self-encoded proteins via genetic variation / evolution to counteract host multi-layered defensive responses, including but not limited to RNA silencing and cellular autophagy. The 6K1 is one of the most evolutionarily-conserved proteins among potyviruses, but its biological relevance is less annotated. This study demonstrates that most of conserved residues in potyviral 6K1s are essential for a successful infection of PVMV. We provide multi-disciplinary evidence supporting that cellular autophagy is engaged in the degradation of 6K1. We defined a conserved lysine/arginine-rich motif in 6K1s across potyviruses that is responsible for the autophagy-mediated self-degradation to promote viral infection. This finding provides a new insight in our understanding of a conserved but understudied potyviral protein.

Numerous studies demonstrate that autophagy plays dual roles during virus-plant interactions (36–38). Regarding potyviruses, autophagy-mediated degradation of key viral proteins such as HCPro and NIb fight against viral infection (40–42). This study proves that autophagy-mediated degradation of 6K1 facilitates viral infection, indicating a pro-viral role of autophagy in potyviral infection. Previously, there were two reports showing the facilitative effect on potyviral infectivity: ⅰ) TuMV VPg interacts with SGS3 to mediate the autophagic degradation of both SGS3 and RDR6 to promote viral infection (43); ⅱ) TuMV P1 protein interacts with cpSRP54 to mediate its autophagic degradation to suppress JA biosynthesis (46). The two cases reveal that potyviral proteins cooperate or co-opt cellular autophagy machinery to eliminate core proteins in antiviral pathways. Based on these observations, we envisage a scenario that 6K1 cooperate with autophagy machinery to degrade certain cellular antiviral component. To verify this hypothesis, two following questions need to be answered.

First, what is the molecular mechanism underpinning the autophagic degradation of 6K1? To address this question, it is necessary to perform a comprehensive screening of potential interactors of 6K1 to identify autophagic receptor and adaptor proteins, which will be a promising research direction. Recently, a milestone discovery for potyvirid 6K1 is that it is also an endoplasmic reticulum (ER)-localized integral membrane protein, forms pentamers with a central hydrophobic tunnel, and increases the cell membrane permeability to facilitate viral infection (58). Intriguingly, artificial intelligence-assisted structure modeling and biochemical assays demonstrated that three arginine residues, i.e., K/R34, K38, and K40 in the conserved Lys/Arg-rich motif defined in this study (Fig. 2), are responsible for its oligomerization (58). This prompted us to speculate that the four residues (V32, K34, L36, and L39) in Lys/Arg-rich motif that determine 6K1’s autophagic degradation, might depend on its oligomerization. Our study emphasizes the importance of the conserved Lys/Arg-rich motif in 6K1’s self-interaction for implementing critical biological functions.

Second, which components are co-degraded with 6K1 by autophagy. Interestingly, a recent study showed that transient expression of TuMV 6K1 decreases the activity of cellular cysteine proteases (59), which were demonstrated as the central hubs of plant immunity against many pathogens including viruses (60–63). SMV 6K1 interacts with a large number of defense-related proteins from soybean, such as pathogenesis-related protein 4, Bax inhibitor 1, papain family cysteine protease, and cysteine protease inhibitors (53). PVY 6K1 interacts with 14-3-3 protein, playing a vital role in plant defense against various pathogens, including PVY (32). It seems that the known interactors of 6K1 are all categorized as defense-related proteins. Consequently, the clarification of host components co-degraded with 6K1 would unveil a novel counter-defense strategy expressed by potyviruses.

In the case of PPV, artificial inactivation of proteolytic processing at P3-6K1 junction via mutating either ‘Gln to His’ at P1 position or ‘Ala to Lys’ at P1′ position, markedly weakened viral systemic infection in *N. benthamiana*, suggesting an accessory role of 6K1 in viral multiplication (25, 26, 30). However, the effects of introduced mutations on NIa-Pro processing of P3-6K1 junction were detected in *in vitro* assays. The influences of these mutations on the processing *in planta* await to be proved. For PVMV, these mutations, when introduced into viral clones, abolished the P3-6K1 processing *in planta*, leading to a failure of viral infection. Spontaneous reversion mutation from Lys to Asn at P1′ position restored the proteolytic processing, and thus rescued viral systemic infection. These results strongly suggest that proteolytic processing at P3-6K1 junction to release mature 6K1 is required for viral successful infection. In addition, we observed that a slow processing at P3-6K1 junction had no obvious effect on viral systemic infection and symptom development (Fig. 4). Nevertheless, the correlation between the processing of P3-6K1 junction *in planta* and viral infectivity needs to be carefully examined in different potyvirus-host pathosystems.

Plant viruses characteristically have small sizes and compact genomes. To overcome the limited coding capacity, viruses evolved a variety of strategies to express more viral proteins, such as stop codon readthrough, leaky scanning, and frameshifting. In *in vitro* assays, both the cleavages at HCPro-P3 junction by HC-Pro and at 6K1-CI junction by NIa-Pro is efficient (27, 28, 64, 65), whereas the cleavage at P3-6K1 junction is slow (25, 27), suggesting the generation of three viral proteins (i.e., P3, 6K1, and P3-6K1) from corresponding genomic region (26). Consistently, all PVMV-derived clones tested in this study, if variable, expressed the precursor P3-6K1 to a varied degree (Figs. 3E, 3F, 4D, 4E). Based on these observations, we assume that: ⅰ) P3-6K1 might be also functional, which deserves further investigation. In the case of TuMV, the precursor P3-6K1 is an integral membrane protein, whereas P3 is a peripheral membrane protein; P3-6K1 forms small granules on the ER network, and might have distinct biological function(s) (58). ⅱ) besides polyprotein processing and RNA polymerase slippage, the incomplete processing at intercistronic junctions is an alternative expression strategy for potyvirids, which deserve being paid more attention in future. Potyviral 6K1s are short in size (∼54-aa), and 17 residues are rather conserved. It is so fascinating that 15 of them are essential for the successful infection of PVMV. Are these residues responsible for the functions of 6K1, P3-6K1, or both? To best define which residues are associated with biological significance of 6K1, these 6K1 variants with single alanine/arginine substitution of conserved residues are individually engineered into viral clone, and allowed for their expression along with viral infection, in order to interfere with the functions of inherent 6K1 in PVMV. Fortunately, four point mutants interfere with viral infectivity, but also are associated with autophagic degradation of 6K1. In view of the limitations of this approach, it would be also possible that the remaining conserved residues involve biological function of 6K1, could not be precluded, or else, some residues are associated with the potential functions of P3-6K1. For potyvirids, viral factors excluding P3N-PIPO are expressed through polyprotein processing, and theoretically, should share an equivalent number of molecules during viral infection. In fact, an extremely low abundance of 6K1 was observed in the context of viral infection for all tested potyviruses (28, 30). In line with the observation, we provide multi-disciplinary evidences in supporting a role of cellular autophagy on the degradation of PVMV 6K1. Curiously, E-64d treatment increased the accumulation level of 6K1, whereas 3-MA does not. The contrasting effects (3-MA versus E-64d treatments) might be well explained by following facts: ⅰ) 3-MA is able to effectively block an early stage of autophagy by inhibiting the formation of class III PI3K complex, but does not inhibit Beclin1/VPS30/ATG6-independent autophagy pathways (66–68). Beclin1 is a core component in the formation of autophagosomes. However, in mammalian cells, it has been shown that several types of autophagy are induced in a Beclin1/VPS30/ATG6-independent manner, and are not blocked by PI3K inhibitors (66, 69, 70). The formation of autophagosomes in this functional autophagy pathway just need a subset of ATG proteins (66, 71). ⅱ) Several studies showed that 3-MA might also stimulate autophagy (72, 73), suggesting that it is not a preferable inhibitor for autophagy-related research. ⅲ) E-64d blocks the late steps of autophagy pathways through inhibiting the activity of aspartic and cysteine proteases (74). Our finding increases the possibility that Beclin1/VPS30/ATG6-independent autophagy pathway might occur in plants.

## MATERIALS AND METHODS

### Plant materials and virus source

*N. benthamiana* and *C. chinense* (cultivar Yellow Lantern) plants were grown in a growth chamber with the set of growth conditions as follows: 16-h light (6500 Lx) at 25°C and 8-h darkness at 23°C with the relative humidity of 70%. The infectious cDNA clone of a PVMV isolate (PVMV-HNu), pHNu, was developed by our group (48), and served as the backbone to produce a series of indicated recombinant and mutated viral clones.

### Construction of PVMV-derived cDNA clones

To create a GFP-tagged PVMV clone, the NIb/CP intercistronic junction in pHNu (48) was subjected for the integration of a complete GFP-encoding sequence. The GFP sequence was amplified from pVPH-GFP (30) with a pair of primers P-GFP-F/P-GFP-R (Table S1). Another two regions, upstream and downstream of NIb/CP junction, respectively, were amplified with corresponding primer sets C-F/C1-R and C2-F/C-R (Table S1). A mixture of above obtained products was used as the template for overlapping PCR with primer set C-F/C-R. The resulting fragment was inserted back into pHNu (48) by utility of *Aat*Ⅱ/*Sal*I sites to generate the recombinant clone, pHNu-GFP.

To engineer a second copy of 6K1 into NIb/CP junction, the 6K1-coding sequence was amplified by primer set NIb-6K1-F/6K1-CP-R (Table S1). Another two regions, upstream and downstream of NIb/CP junction, respectively, were amplified with corresponding primer sets C-F/NIb-6K1-R and 6K1-CP-F/C-R (Table S1). A mixture of the obtained PCR products was used as the template for overlapping PCR with the primer sets C-F/C-R. The resulting fragment was inserted into pHNu by utility of *Aat*Ⅱ/*Sal*I sites to generate the clone pHNu//6K1. Using pHNu//6K1 as the backbone, a total of 15 mutated clones, i.e., pHNu//6K1(K3A), pHNu//6K1(E11A), pHNu//6K1(A15R), pHNu//6K1(L19A), pHNu//6K1(M22A), pHNu//6K1(D25A), pHNu//6K1(D27A), pHNu//6K1(R28A), pHNu//6K1(R30A), pHNu//6K1(V32A), pHNu//6K1(K34A), pHNu//6K1(L36A), pHNu//6K1(K38A), pHNu//6K1(L39A), and pHNu//6K1(K51A), were constructed for the generation of recombinant viruses with the mutation of corresponding conserved residues in the second copy of 6K1 into Ala or Arg. These clones were generated in the same strategy, and, herein, we described the construction of pHNu//6K1(K3A). Two primers K3A-F and K3A-R (Table S1) were synthesized, and, respectively, paired with C-R and C-F for PCR reactions with pHNu//6K1 as the template. A mixture of the obtained PCR products was used as the template for overlapping PCR with the primer sets C-F/C-R. The resulting fragment was inserted into pHNu//6K1 by utility of *Aat*Ⅱ/*Sal*I sites to generate the clone pHNu//6K1(K3A).

To integrate a GFP-tagged 6K1 into NIb/CP junction, one upstream fragment of 6K1-CP junction in pHNu//6K1 was amplified using primer set C-F/6K1-GFP-R (Table S1), another downstream fragment of NIb-GFP junction in pHNu-GFP was amplified with primer set 6K1-GFP-F/C-R (Table S1). Two PCR products were mixed, and served as the template for overlapping PCR with primer set C-F/C-R. The resulting fragment was inserted into pHNu by utility of *Aat*Ⅱ/*Sal*I sites to generate pHNu//6K1-GFP. Using a similar cloning strategy for the generation of pHNu//6K1(K3A), the following mutated clones based on pHNu//6K1-GFP were generated: pHNu//6K1(D30A)-GFP, pHNu//6K1(V32A)-GFP, pHNu//6K1(K34A)-GFP, pHNu//6K1(L36A)-GFP, and pHNu//6K1(L39A)-GFP.

To C-terminally fuse a Myc epitope with 6K1 protein in PVMV, two PCR reactions with pHNu-GFP as the template were performed by using primer sets 2280-F/6K1^Myc^-R and 6K1^Myc^-F/4070-R, respectively (Table S1). The obtained products were mixed and served as the template for overlapping PCR with primer set 2280-F/4070-R. The resulting fragment was inserted back into pHNu-GFP by utility of *Bam*HI/*Stu*I sites to generate the clone, pHNu-GFP-6K1^Myc^. pHNu-GFP-^Myc^P3 was constructed by using a similar strategy with that of pHNu-GFP-6K1^Myc^. To destroy the cleavage at P3-6K1 junction by NIa-Pro, five mutated clones, i.e., pHNu-GFP-^Myc^P3(Q-H), pHNu-GFP-^Myc^P3(A-K), pHNu-GFP-^Myc^P3(QA-AQ), pHNu-GFP-^Myc^P3(Q-A), and pHNu-GFP-^Myc^P3(A-Q), were constructed with pHNu-GFP-^Myc^P3 as the backbone using a similar cloning strategy. Here, we described the construction of pHNu-GFP-^Myc^P3(Q-H). Two primers Q-H-F and Q-H-R (Table S1), for which the original nucleotides coding for Gln at P1 position were substituted for encoding His, were synthesized, and, respectively, paired with 4070-R and 2280-F (Table S1) for PCR reactions. A mixture of obtained products served as template for overlapping PCR with primer set 4070-R and 2280-F. The resulting fragment was inserted back into pHNu-GFP-^Myc^P3 by utility of *Bam*HI/*Stu*I sites. Similarly, a total of 17 point mutants of PVMV-GFP-^Myc^P3, in which the conserved residues in 6K1 were substituted with Ala or Arg, were developed.

### Generation of T-DNA constructs

A binary plant-expression vector - pCaMterX (75) was employed to transiently express genes of interest in *N. benthamiana*. The coding sequence of GFP and mCherry were amplified from pVPH-GFP//mCherry (30) using primer sets GFP-F/GFP-R and mCherry-F/ mCherry-R (Table S1), respectively. The resulting fragments were individually integrated into pCaMterX by utility of *Xba*Ⅰ/*Bam*HⅠ sites to generate the clones, pCaM-GFP and pCaM-mCherry. To develop the constructs for transiently expressing NbATG8a and NbATG8f whose N-termini were fused with mCherry, the coding sequences of NbATG8a and NbATG8f were amplified by using cDNAs prepared from *N.benthamiana* leaves with primer sets NbATG8a-Bam-F/NbATG8a-Kpn-R and NbATG8f-Bam-F/NbATG8f-Kpn-R (Table S1), respectively. The resulting PCR products were individually integrated into pCaM-mCherry by utility of *Bam*HⅠ/*Kpn*Ⅰ to generate the clones, pCaM-mCherry-NbATG8a and pCaM-mCherry-NbATG8f. The coding sequence of 6K2 was amplified from pHNu-GFP with a pair of primers 6K2-Xho-F/6K2-Xba-R (Table S1), the obtained PCR products were integrated into pCaM-mCherry by utility of *Xho*Ⅰ/*Xba*Ⅰ sites to generate the clone, pCaM-6K2-mCherry. Similarly, the construct pCaM-6K1-GFP for transiently expressing 6K1-GFP was generated. For the hairpin-mediated silencing in *N. benthamiana*, a partial sequence of coding region of *NbATG7* (324 nt) or β-glucuronidase gene (310 nt) was cloned into p2300s-intron in both sense (*Sac*Ⅰ/*Bam*HⅠ) and antisense (*Pst*Ⅰ/*Xba*Ⅰ) orientations. All PCRs were performed with Phusion high-fidelity DNA polymerase (Thermo Fisher Scientific), and all constructs were confirmed by sanger sequencing.

### Sequence analysis

A total of 115 sequences of different potyviral 6K1s (Supplemental Data S1) were retrieved from NCBI GenBank database, and subjected to multiple alignment. Multiple alignment was performed using an online tool Clustal Omega (https://www.ebi.ac.uk/Tools/msa/clustalo/) (76), followed by the description with the online program Weblogo (http://weblogo.berkeley.edu/logo.cgi) (77, 78).

### Viral inoculation and agroinfiltration

For infectivity test of PVMV-derived cDNA clones, agrobacterium (strain GV3101) cultures harboring corresponding clones were suspended in infiltration buffer (10 mM MgCl_2_, 10 mM MES, and 150 μM acetosyringone), adjusted to OD_600_ of 1.0 and infiltrated into fully expanded leaves of *N. benthamiana* seedlings at 6- to 8-leaf stage or *C. chinense* plants at 3- to 4-leaf stage, unless otherwise stated. For subcellular localization assays, agrobacterium (GV3101) culture harboring pHNu//6K1-GFP (OD_600_, 0.5), was mixed with another culture harboring either pCaM-mCherry-NbATG8a or pCaM-mCherry-NbATG8f (OD_600_, 0.5) in equal proportions, and co-infiltrated into fully expanded leaves of *N. benthamiana* plants at 6- to 8-leaf stage.

To test whether 6K1 is colocalized with 6K2, fully expanded leaves of *N. benthamiana* were firstly infiltrated with agrobacterial culture harboring pHNu//6K1-GFP (OD600, 0.5). 24 h later, these patches were re-infiltrated with an agrobacterial culture harboring pCaM-6K2-mCherry (OD_600_, 0.3). For hairpin-mediated silencing assays, each of agrobacterium cultures harboring pCaM-6K1-GFP or its mutants, together with another culture containing p2300s-intron-dsGUS or p2300s-intron-dsATG7, were adjusted to OD_600_ of 1.0, mixed with equal volumes, and co-infiltrated into fully expanded leaves of *N. benthamiana* plants at 6- to 8-leaf stage.

### RNA analysis

Total RNAs were extracted from leaf tissues of *N. benthamiana* with TRNzol reagent (TIANGEN) and *C. chinense* with FastPure Plant Total RNA Isolation Kit (Vazyme). For RT-qPCR analysis, total RNAs (1 μg per sample) were treated with DNaseI (Thermo Fisher Scientific) following the manufacturer’s instructions, followed by reverse-transcription with RevertAid First Strand cDNA Synthesis Kit (Thermo Fisher Scientific). The synthesized cDNAs were used to determine mRNA levels of target genes or quantification of viral accumulation levels. Specific primer pairs were designed using Primer3Plus (https://www.primer3plus.com/index.html) (79). qPCR was conducted by using SuperReal Premix Plus (TIANGEN) in Applied Biosystems QuantStudio 5 (Thermo Fisher Scientific). The transcripts of either *NbActin* or *CcActin* were selected as an internal control to normalize the data. Each experiment was performed at least three times, and the relative gene expression level were calculated by manufacturer’s software.

### Immunoblotting and antibodies

Total proteins were extracted from infiltrated leaf patches or systemically infected leaves of *N. benthamiana* and *C. Chinense*, following a previously described protocal (30). Immunoblotting assays were performed with anti-GFP rabbit polyclonal antibody (BBI) or anti-Myc polyclonal antibody (Abcam) as the primary antiboty and horseradish peroxidase (HRP)-conjugated goat anti-rabbit IgG (BBI) or goat anti-rabbit immunoglobulin antibody (Abcam) as the secondary one, essentialy as described previously (80). The immunological detection of target signals was performed using enhanced chemiluminescence detection reagents (Thermo Fisher Scientific) in an ImageQuant LAS 4000 mini biomolecular imager (GE Healthcare). The signal intensity corresponding to protein bands was quantitatively analyzed with ImageJ software (81).

### Chemical treatments

Fully expanded leaves of *N. benthamiana* plants, pre-inoculated with the indicated plasmids, were treated with 1% DMSO in phosphate buffer (as the parallel control), or an equal volume buffer containing 1% DMSO and 10 mM 3-MA, 100 μM E64d or 100 μM MG132 (Sigma). Sixteen hours after the treatment, leaf samples were collected for immunoblotting analysis.

### Confocal microscopy

The epidermal cells of infiltrated patches were examined by a confocal microscopy (LSM 900, Zeiss) with a 20× dry immersion objective. Light emitted at 643 nm was used to record chlorophyll auto-fluorescence; GFP was excited at 493 nm and captured at 478-535 nm; mCherry was excited at 553 nm and captured at 548-629 nm. Images were captured digitally and handled using Zeiss ZEN 3.7 software.

### Accession numbers in NCBI GenBank database

PVMV-HNu (MN082715), *NbATG8a* (KX120976), *NbATG8f* (KU561372), *NbATG7* (KX369398), *CcActin* (AM168448).

## ACKNOWLEDGMENTS

This work is supported by grants from the National Natural Science Foundation of China (31860487), and Collaborative Innovation Center of Nanfan and High-Efficiency Tropical Agriculture, Hainan University (XTCX2022NYB11). We would like to express our special appreciation to Dr. Yi Xu (Nanjing Agricultural University) for providing the plasmid p2300s-intron, and Dr. Wenping Qiu for critical reading. H.C., W.H., Z.D., and X.X. conceived and designed the project. W.H., C.D., and L.Q. carried out experiments. H.C. supervised the work. All authors analyzed and discussed the data. H.C., and W.H. wrote the manuscript. All authors reviewed and approved the manuscript. We declare no conflicts of interest.

